# The Second Mitochondrial Activator of Caspases (SMAC) regulates growth, inflammation and mitochondrial integrity in cancer cells

**DOI:** 10.1101/2025.04.03.646995

**Authors:** Tarek Amer, Aladin Haimovici, Susanne Kirschnek, Juliane Vier, Abdul Moeed, Uzochukwu Ukachukwu, Martin Helmstädter, Severine Kayser, Rupert Öllinger, Roland Rad, Arnim Weber, Mohamed Tarek Badr, Georg Häcker

**Affiliations:** Institute of Medical Microbiology and Hygiene, Medical Center, Faculty of Medicine, University of Freiburg, Freiburg, Germany; EMcore, Internal Medicine IV, Faculty of Medicine, Medical Center, University of Freiburg, Freiburg, Germany; Institute of Molecular Oncology and Functional Genomics, Department of Medicine II and TranslaTUM Cancer Center; TUM School of Medicine, Technical University of Munich, Munich, Germany; Institute of Medical Microbiology and Hygiene, Medical Center, University of Freiburg, Faculty of Medicine, Freiburg, Germany; Signaling research centres BIOSS and CIBSS, University of Freiburg, Freiburg, Germany

## Abstract

SMAC is a mitochondrial intermembrane space protein, which is released during apoptosis and whose known function is antagonism of inhibitor of apoptosis proteins in the cytosol, to facilitate caspase activation. Recent data suggest that SMAC can also be released by sub-lethal signals in the apoptosis pathway, in the absence of cell death. We here explored potential functions of SMAC in non-apoptotic cells. We found that a portion of SMAC is spontaneously released into the cytosol in the absence of apoptosis, regulated by the BCL-2-family proteins BAX and BAK and the fission GTPase DRP1. In cancer cell lines, SMAC was required for the activation of caspases in lethal and non-lethal conditions, while this contribution to caspase-activation was much smaller in non-malignant fibroblast lines. In cells with high levels of cytosolic SMAC, SMAC deficiency reduced *in vitro* migration, invasion and anchorage-independent growth. SMAC-deficient cells further showed a reduced activity in interferon signalling, associated with reduced cytosolic presence of mitochondrial DNA and activation of the stimulator of interferon genes (STING), and SMAC expression levels correlated with interferon-induced genes in cancer data sets. We further found that SMAC can regulate mitochondrial morphology and integrity. Finally, high gene-expression of SMAC was associated with poor prognosis in patients of several cancer types. These results identify SMAC as a regulator of inflammation and growth behaviour of cancer cells. They further report a mitochondrial function of SMAC and demonstrate a role of SMAC in human cancer biology across several cancer entities.

## Introduction

One of the functions of mitochondria is the regulation of apoptosis ^1^. Following the integration of upstream signals, BCL-2-family proteins can permeabilize the outer mitochondrial membrane and release intermembrane space proteins that trigger the cytosolic activation of caspases ^2^. Outer membrane permeabilization eventually kills the cell; caspase-activation causes the apoptotic morphology and counteracts the inflammatory activity of mitochondrial contents such as mitochondrial (mt) DNA ^3^.

The release of cytochrome *c* is indispensable for caspase-activation and apoptosis ^4^. SMAC, also known as Direct IAP Binding protein with Low PI (DIABLO), was identified as an inhibitor of IAPs (inhibitor of apoptosis proteins) in the cytosol ^5, 6^. SMAC translocates to the cytosol alongside cytochrome *c* during apoptosis ^7^ and in the cytosol releases the inhibition of XIAP on caspase-9 ^8^. In a number of experimental models, for instance in TRAIL signalling, this XIAP inhibition has been found to be required for apoptosis ^9^, and the role of the SMAC-XIAP interaction for apoptosis has been modelled mathematically ^10, 11^. However, SMAC appears to be variably relevant for apoptosis depending on cell type. Analysis of SMAC-deficient mice suggests no or little importance of SMAC in thymocyte, lymphocyte, embryonic stem cell or fibroblast apoptosis ^12^. On the other hand, SMAC was essential for the activation of caspases upon mitochondrial permeabilization in human cell lines ^13^. Small molecules have been developed that mimic the molecular function of SMAC ^14, 15^. These molecules have identified a potential pro-inflammatory role of SMAC by antagonism of cIAP1/2 ^16^.

SMAC is a mitochondrial protein: like most mitochondrial proteins it is encoded in the nuclear genome, synthesized in the cytosol and imported into mitochondria. During import, its N-terminus is cleaved by the protease PARL ^13^. SMAC has also been proposed to regulate phospholipid synthesis during tumorigenesis ^17^, and A549 lung cancer cells deficient in SMAC showed reduced growth and tumorigenesis ^18^. SMAC may therefore have functions in non-apoptotic cells ^19^ but it is unclear how it achieves these effects.

During apoptosis, the release of cytochrome *c* into the cytosol is rapid and complete ^20^, and the kinetics of SMAC release follow a similar pattern ^7^. Recent results show that the release of mitochondrial cytochrome *c* can be partial when cells are treated with a sub-lethal dose of BCL-2 inhibitor. In this situation, only a small portion of mitochondria permeabilize, and the level of caspase-activation elicited is too low to kill the cell (minority MOMP) ^21, 22^. Such sub-lethal signalling has been found in cellular infection with numerous pathogens ^23^, and during infection with the bacterium *Helicobacter pylori*, SMAC can be released in the absence of cell death but in sufficient amounts to cause NF-κB-activation ^24^. SMAC may therefore have a function in sub-lethal signalling following its release into the cytosol. It is further surprising that SMAC is a mitochondrial protein but its reported molecular functions are exclusively cytosolic. Lastly, an unexpected correlation of SMAC-expression and tumour progression has been reported. Given its function as a pro-apoptotic protein and the general role of apoptosis as a cancer-preventing mechanism, a correlation of high levels of SMAC and a good prognosis in cancer may be expected. However, the opposite was the case in metastatic melanoma: high SMAC-levels were linked to poor survival ^25^.

We here tested whether non-lethal release of SMAC into the cytosol may be relevant for its biology. We observed spontaneous cytosolic presence of SMAC in human cell lines, which was relevant for the regulation of growth behaviour. SMAC levels further regulated the interferon system in cancer cells, associated with the cytosolic presence of mtDNA. SMAC-deficient cells showed changes in their mitochondrial morphology and integrity.

## Results

### Requirement for SMAC in lethal and sub-lethal caspase-activation

As discussed above, the dependency of cells on SMAC for caspase-activation following mitochondrial permeabilization appears to be variable. We deleted SMAC in three human cancer cells lines (HeLa cervical carcinoma, MDA-MB231 breast cancer and 1205Lu melanoma cells) and tested for the requirement for SMAC in caspase-activation (gene deletions for all cell lines used here are documented in Fig. S1). The activation of caspase-3 during induction of apoptosis with the combination of the BCL-2/BCL-X_L_-inhibitor ABT-737 and the MCL-1-inhibitor S63845 was strongly SMAC-dependent in these cell lines (Fig. 1A), expanding earlier results ^13^. This clear SMAC dependency is in obvious contrast to the findings in primary cells from SMAC-deficient mice where no apoptosis defect had been observed ^12^. We therefore tested non-transformed cells from mouse and human origin. SMAC-deficient human WI-38 fibroblasts and two mouse embryonic fibroblast lines established from SMAC deficient mice did show some reduction in apoptosis sensitivity to BH3-mimetics but this effect was relatively subtle (Fig. 1B). There were no obvious differences in XIAP expression that could have explained this strong difference (Fig. S2). With this sample of cells, it appears that cancer cell lines show a substantially higher dependency than non-malignant cells on SMAC for caspase-activation downstream of cytochrome *c*-release.

**Figure 1:**
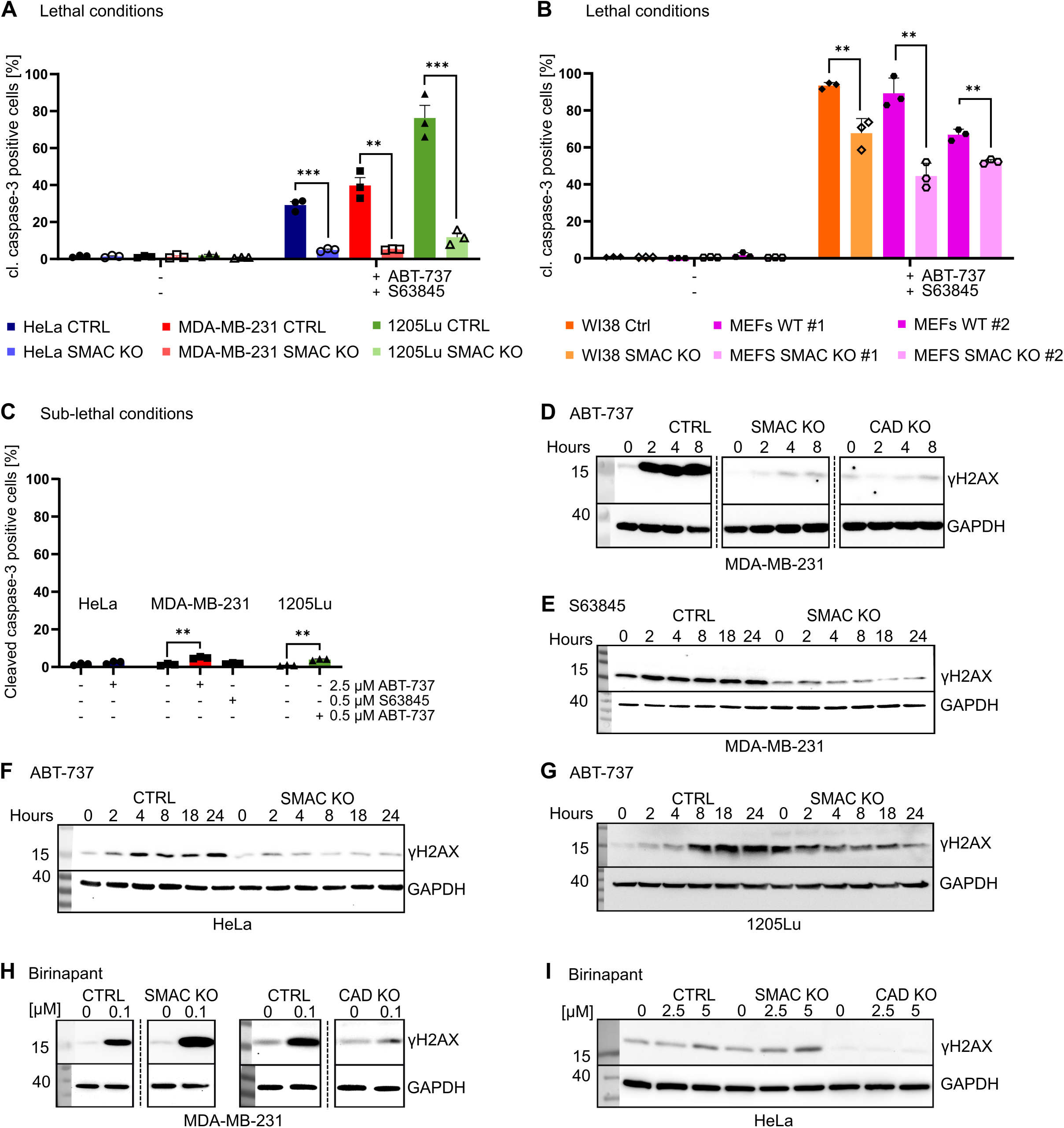
SMAC is required for lethal and sub-lethal mitochondrial signaling in cancer cell lines. **A**, the human cancer cell-lines HeLa (cervical carcinoma), MDA-MB-231 (breast carcinoma) and 1205Lu (metastatic melanoma), each control (CTRL; carrying a non-targeting gRNA) or SMAC-deficient or **B**, WI38 human fibroblasts WI38 (control or SMAC-deficient) or mouse embryonic fibroblasts (established from littermate SMAC deficient or wt mice) were treated with ABT-737 (2.5 µM) and S63845 for 4 h. We used S63845 at 0.5 µM S63845 for human and 5 µM for mouse cells. **C**, HeLa cells, MDA-MB-231 or 1205Lu cells were treated with the inhibitors as indicated (HeLa cells, 4 h with 2.5 µM ABT-737; MDA-MB-231, 2.5 µM for 4 h or 0.5 µM S63845 for 24 h; 1205Lu cells, 0.5 µM ABT-737 for 24 h) (‘sub-lethal’ conditions). Cells were fixed, stained for cleaved caspase-3 and analyzed by flow cytometry. Data are means/SEM of at least three independent experiments. *, p<0.05, **, p<0.01, ***, p<0.001 (unpaired t-test). **D**-**I**, induction of the DNA-damage response marker γH2AX. **D**, MDA-MB-231 CTRL, SMAC-deficient and CAD-deficient cells treated with ABT-737 (2.5□µM) or **E** S63845 (0.5 µM) for the indicated times. **F**, HeLa CTRL and SMAC-deficient cells treated with 2.5 µM ABT-737 for the indicated times. **G**, 1205Lu CTRL and SMAC-deficient cells treated with 0.5 µM ABT-737 for the indicated times. **H**, MDA-MB-231 or **I** HeLa CTRL, SMAC-deficient and CAD-deficient cells were treated with the SMAC mimetic Birinapant for 24 h at the concentrations indicated. Western blots are representative of at least three independent experiments. Dashed lines indicate rearranged lanes from the same membrane.

We established conditions where no or very little cell death was induced (‘sub-lethal conditions’) using low concentrations of either BH3-mimetic (Fig. 1C). We measured sub-lethal signals by the caspase-activated DNase (CAD)-dependent appearance of the DNA-damage response signal γH2AX ^25^. Again, the signal required SMAC in MDA-MB-231 (Fig. 1D, E), HeLa (Fig. 1F) and 1205Lu cells (Fig. 1G). The required activity of SMAC was the inactivation of IAPs, presumably XIAP, because the addition of the small molecule SMAC mimetic/IAP inhibitor Birinapant reversed the effect of SMAC-loss and permitted sub-lethal caspase-activation, as measured by the γH2AX-signal in MDA-MD-231 (Fig. 1H) and HeLa cells (Fig. 1I).

We have previously reported spontaneous activity of the mitochondrial apoptosis pathway in proliferating cells in culture ^25^. Spontaneous caspase-3-activation also depended on SMAC: Western blotting detected cleaved caspase-3 in HeLa and MDA-MB-231 cells, and this required the genomic presence of SMAC (Fig. 2A). Effector caspase (DEVD-cleaving) activity was also detectable, and this activity was substantially reduced in the absence of SMAC in either cell line (Fig. 2B). Active caspase-3 could further be precipitated from control but not from SMAC- or caspase-9-deficient HeLa or MDA-MB231 cells when a biotinylated caspase-substrate was added to the cells prior to cell lysis and processing (Fig. 2C). The results identify the requirement for SMAC in the activation of caspases in the mitochondrial pathway during lethal and sub-lethal signaling. Intriguingly, such requirement was even seen in the absence of any pro-apoptotic stimulus. This suggests that some SMAC is constantly released into the cytosol in steady state to trigger caspase-activation.

**Figure 2:**
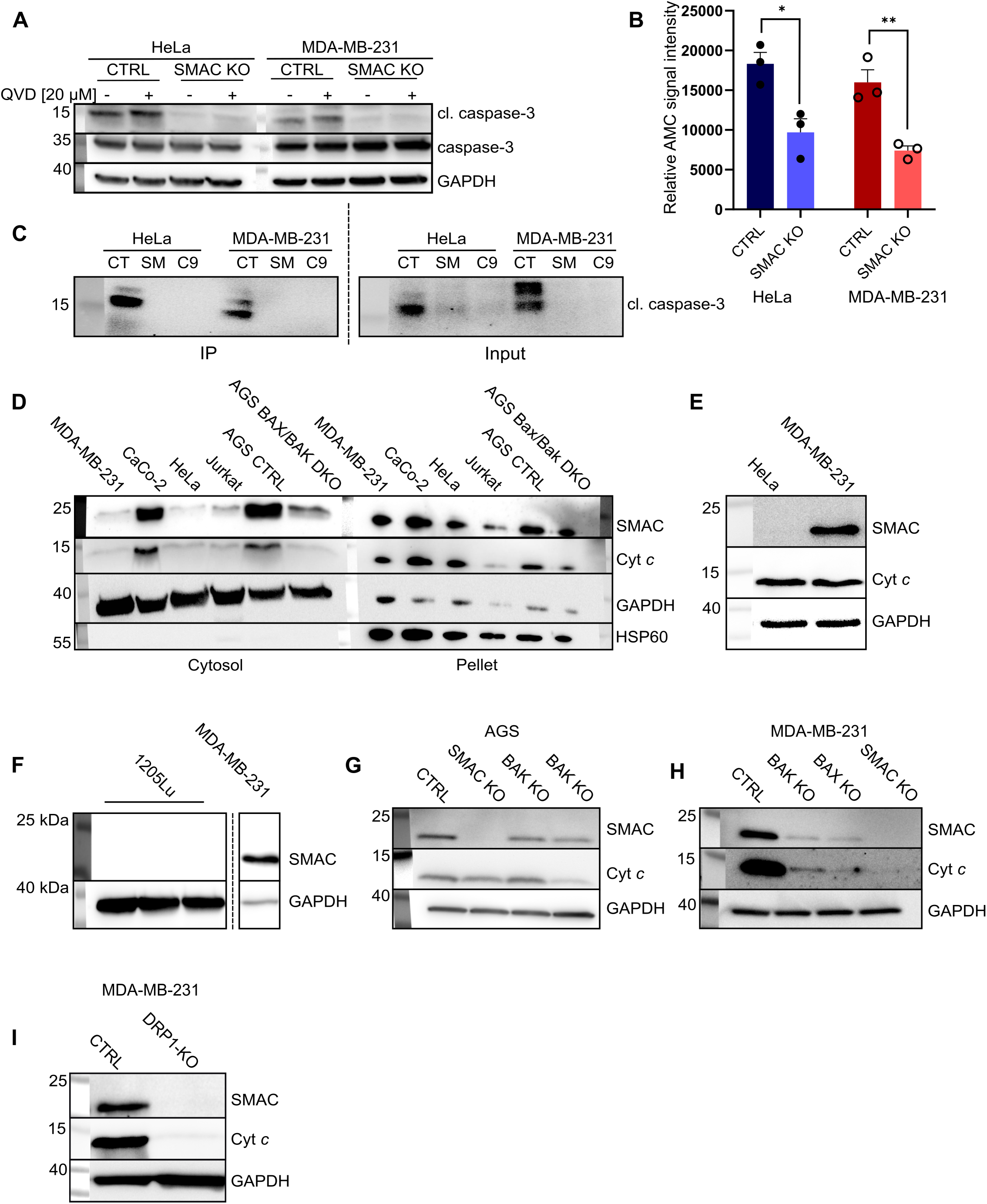
A stable cytosolic fraction of SMAC drives constitutive activation of caspase-3. **A**, HeLa or MDA-MB-231 cells (CTRL and SMAC-deficient) treated for 24 h with the caspase inhibitor QVD as indicated and probed for cleaved caspase-3. **B**, HeLa or MDA-MB-231 (CTRL and SMAC-deficient) cells were analyzed for active caspase-3/-7 by DEVDase assay. Data means/SEM of at least three independent experiments. *, p<0.05, **, p<0.01 (unpaired t-test). **C**, biotinylated VAD-fmk was added to HeLa or MDA-MB-231 CTRL (CT), SMAC-deficient (SM) or caspase-9-deficient (C9) adherent cells. Inhibitor-bound caspases were precipitated from cell lysates using neutravidin beads (IP). Active caspase3 was detected by Western blotting. **D**, MDA-MB-231, Caco-2 (colorectal cancer), HeLa, Jurkat (T cell lymphoma), AGS (gastric carcinoma) cells, CTRL or (for AGS) BAX/BAK-deficient cells were fractionated into cytosol and pellet containing mitochondria. A small volume of the pellet lysates was loaded a technical control. Fractions were investigated for SMAC by Western blotting. **E**, cytosolic fraction of MDA-MB-231 and HeLa were investigated for SMAC using Western blotting. **F**, SMAC levels in cytosolic fraction of 1205Lu were compared to those in the cytosolic fraction of MDA-MB-231 cells using Western blotting. **G**, cytosolic fractions of AGS CTRL, SMAC, BAK or BAX-, BAK-deficient cells. **H**, MDA-MB-231 CTRL, BAX-, BAK- or SMAC-deficient cells or **I**, fractions of MDA-MB-231 CTRL and DRP1-deficient cells were analyzed for SMAC and cytochrome **c**. The Western blots are representative of three independent experiments. Dashed lines indicate rearranged lanes from the same membrane.

### A portion of SMAC is found in the cytosol in non-apoptotic cells

Fractionation of cells into cytosolic and mitochondrial fractions showed that a portion of SMAC, and also of cytochrome *c*, is detectable in the cytosol in unstimulated cells. A signal for SMAC was detected in the cytosol, and this signal varied greatly between cell lines, from a very strong signal in two cell lines from the gastrointestinal tract (AGS, gastric, and CaCo2, colon) to no detectable signal in the metastatic melanoma cell line 1205Lu (Fig. 2D-F). The cytosolic presence of SMAC was strongly reduced but not abolished by the deletion of BAX and BAK in AGS gastric carcinoma cells (Fig. 2D). Individual deletion of either BAX or BAK had a relatively small effect in AGS (Fig. 2G), and a more pronounced effect in MDA-MB-231 cells (Fig. 2H), suggesting that either protein can contribute to spontaneous release of these proteins into the cytosol. During minority MOMP induced by ABT-737, sub-lethal signals were inhibited by forced mitochondrial fusion ^22^. We therefore deleted the GTPase DRP1, which is required for mitochondrial fission. Loss of DRP1 appeared completely to block the spontaneous release of SMAC and cytochrome *c* (Fig. 2I), indicating that both fission and the BCL-2-family can regulate the release.

### SMAC can regulate growth in cell lines and fibroblasts

Spontaneous activity of the mitochondrial apoptosis pathway can affect growth behavior of human and mouse cells ^25^. Because this spontaneous activity depended on SMAC, we tested the effect of SMAC loss on aspects of growth behavior (please note that not all assays work for all the cell lines; for instance HeLa cells grow in soft agar similar to their growth on plastic support, making it impossible to test specifically for anchorage-independent growth). SMAC-deficient MDA-MB231 cells showed reduced migration, reduced invasion through matrix and reduced colony formation in soft agar (Fig. 3A). MDA-MB-231 have easily detectable SMAC levels in the cytosol. In HeLa cells, where the cytosolic SMAC levels were lower, there was no significant difference in wound healing or invasion by SMAC deletion but a small decrease in proliferation (Fig. 3B). In AGS cells, which again have high levels of cytosolic SMAC, SMAC deficiency was also associated with a reduction of colony growth in soft agar (Fig. 3C), while this was not the case for 1205Lu cells. The small or absent effect of SMAC deletion in HeLa and 1205Lu cells is most likely linked to the small amount of cytosolic SMAC in these cells (see Fig. 2E). The result in the two SMAC-deficient MEF lines was heterogeneous, with one but not the other showing a clear growth deficiency (Fig. 3E). The data suggest that SMAC, which is spontaneously released from mitochondria into the cytosol, affects growth behavior of cancer cell lines.

**Figure 3:**
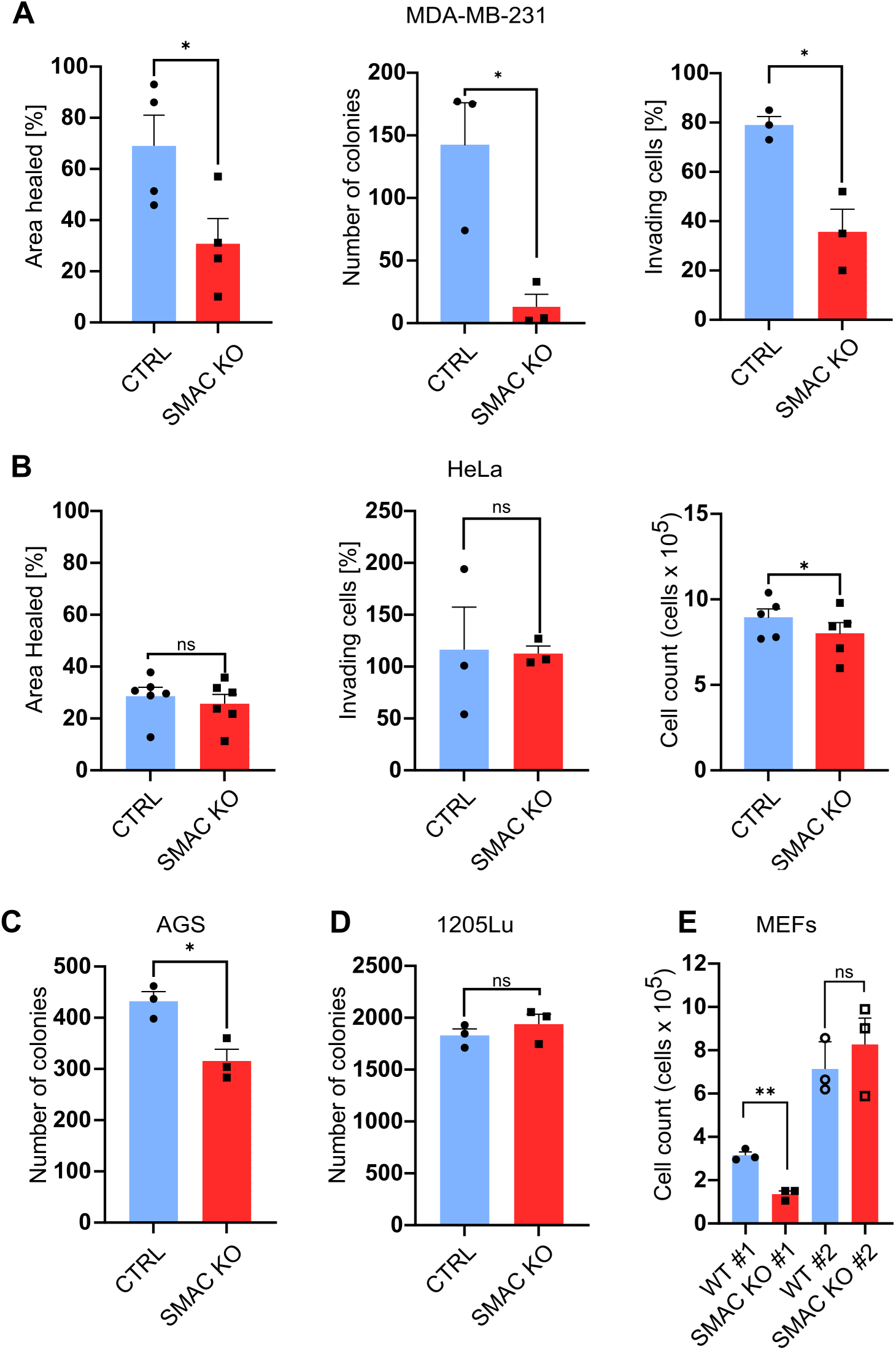
SMAC can regulate cellular growth behavior. CTRL or SMAC-deficient cells from the cell lines MDA-MB-231 (**A**), HeLa (**B**), AGS (**C**) or 1205Lu (**D**) were tested in various assays of growth behavior. Wound healing by scratch assay (**A**, **B**), colony formation in soft agar (**A**, **C**, **D**), invasive growth in matrigel (A) and cell number (**B**) were tested as indicated. **E**, growth of 3T3 immortalized MEFs (wt or SMAC-deficient) was measured by cell counting 48 h after seeding of equal numbers of cells. Means and SEM are shown, symbols give results from individual experiments. ns, p>0.05, *, p<0.05, **, p<0.01 (unpaired t-test). Note that not all cell lines are suited for the various assays.

### SMAC controls interferon response and mtDNA release

These results suggested that SMAC has cytosolic functions beyond the regulation of caspases in apoptosis. We conducted RNA-sequencing in control and SMAC deficient MDA-MB-231 cells. To control for effects due to caspase-activation by SMAC, we included caspase-9-deficient cells. Principal component analysis showed a clear separation of SMAC-deficient cells from control cells while caspase-9 deficiency had only a very small effect (Fig. 4A). Comparing control to SMAC-deficient cells, we identified a set of 1,444 differentially regulated genes, while caspase-9-deficient cells showed a difference from control cells only in 13 genes (10 of which were also differentially regulated in SMAC-deficient cells) (Fig. 4B). Gene enrichment analysis showed a strong downregulation of genes known from the interferon (IFN) response in SMAC-deficient cells compared to control cells (Fig.4C).

**Figure 4:**
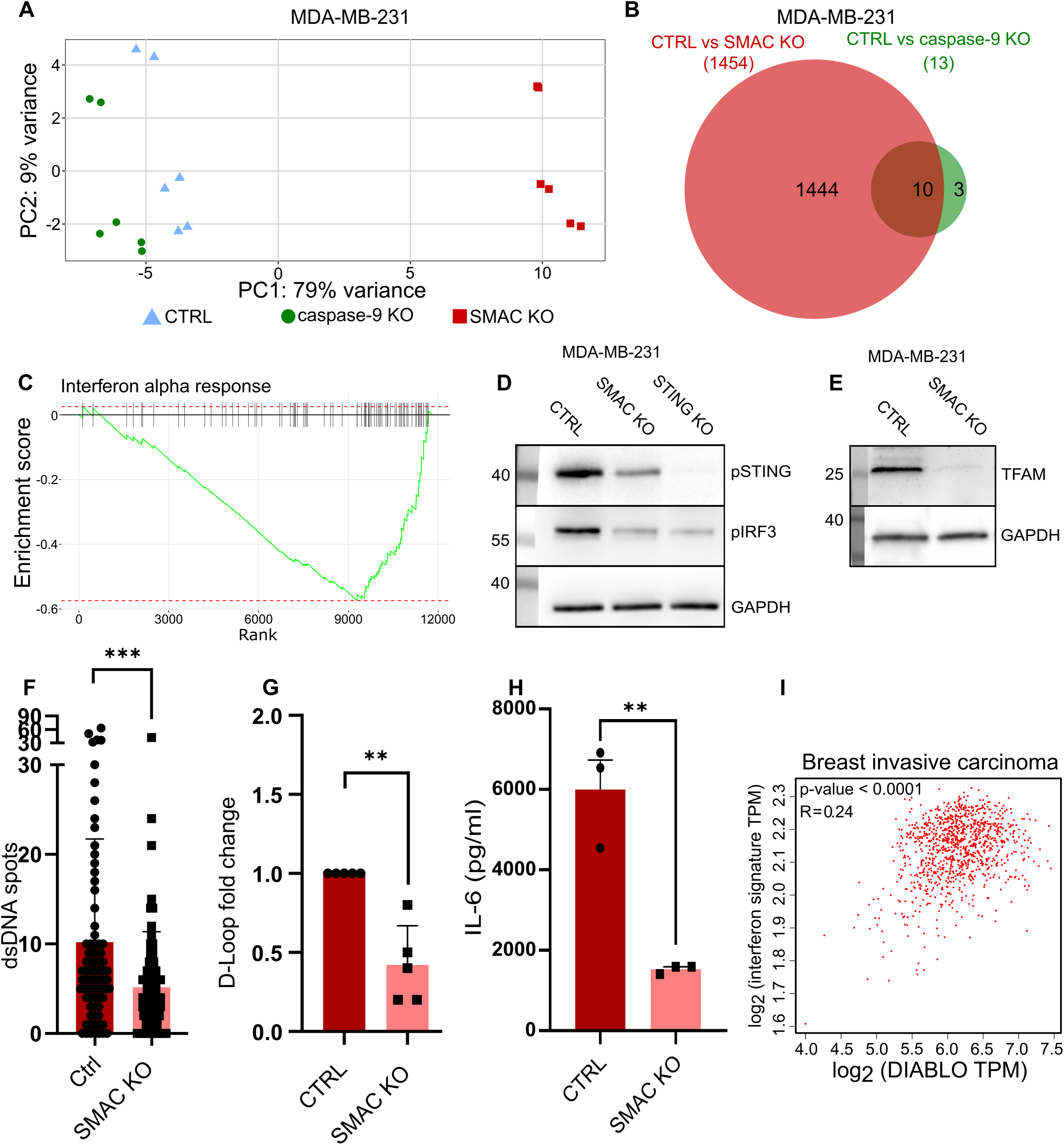
loss of SMAC drives an interferon response characterized by STING phosphorylation and an increase in cytosolic mtDNA in MDA-MB-231 cells. **A**, principal components analysis shows clustering of MDA-MB-231 CTRL and caspase-9-deficient cells while SMAC-deficient cells form a separate cluster. **B**, SMAC-deficient cells have 1,444 differentially expressed genes (q<0.05) while caspase-9-deficient cells show only 13 differentially expressed genes when compared to CTRL cells; ten of these genes are also differentially expressed in SMAC-deficient cells. **C**, gene enrichment analysis shows downregulation of genes associated with interferon alpha response upon loss of SMAC in MDA-MD-231 cells. **D**, levels of phosphorylated STING and IRF3 in CTRL and in SMAC-deficient MDA-MB-231 cells. **E**-**G**, comparison of CTRL and SMAC –deficient MDA-MB-231 cells: **E**, levels of TFAM in the cytosolic fractions, **F**, immunostaining and confocal analysis for dsDNA in the cytosol, **G**, quantification of cytosolic mtDNA using qPCR for a D-loop sequence. Signal detected in cytosol was normalized to the signal detected in whole-cell lysates. **H**, MDA-MB-231 CTRL and SMAC-deficent cells were cultured for 48h. Supernatants were collected and IL-6 was measured by ELISA. Shown are means/SEM of at three experiments. **I**, correlation of the expression levels of the DIABLO gene (encoding SMAC) and interferon-regulated genes in tumor samples from breast invasive carcinoma. Pearson correlation coefficient is shown. **F**, symbols represent individual cells; **G** and **H**, symbols show the results from individual experiments. **I**, each dot represents one tumor sample. Data are means/SEM. **, p<0.01, ***. P<0.001 (unpaired t-test (F and H), Mann-Whitney test (G). The Western blots are representative of at least three independent experiments.

An IFN response is often driven by the stimulator of interferon genes (STING), which signals upon recognition of cytosolic DNA by the receptor of dsDNA, cGAS. We identified easily detectable levels of phorphorylation of STING and of the key transcription factor of the IFN pathway, IRF3, in MDA-MB-231 control cells, which was substantially reduced in SMAC-deficient cells. The levels of IRF3 phosphorylation were comparable in SMAC- and in STING-deficient cells, compatible with the interpretation that SMAC-dependent STING activation accounted for a large part of IRF3 phosphorylation (Fig. 4D). A well-characterized ligand of cGAS is mitochondrial DNA (mtDNA), which can be released into the cytosol ^26^. We found that SMAC-deficient cells had less mtDNA in their cytosol compared to control cells, indicated by reduced cytosolic levels of the mitochondrial transcription factor A (TFAM), reduced cytosolic staining for cytosolic dsDNA and reduced mtDNA in cytosolic fractions of SMAC-deficient cells measured by quantitative PCR (Fig. 4E-G). SMAC-deficient MDA-MB-231 cells further secreted reduced amounts of IL-6, a cytokine that is at least partly regulated by STING ^27^ (Fig. 4H). To test for this SMAC-dependent regulation of the IFN response in cancer, we analyzed gene sets of the TCGA database. In a large number of cancer entities, we detected a significant positive correlation of SMAC expression and the expression of IFN-regulated genes (table S1); Fig. 4I shows the data for invasive carcinoma of the breast. These results suggest the surprising conclusion that the loss of SMAC leads to a reduction of spontaneous release of mitochondrial DNA into the cytosol, and this may account for the correlation of SMAC expression and the IFN response in human cancer.

### SMAC loss affects mitochondrial characteristics

SMAC therefore appears to have control over the release of mtDNA. mtDNA can be released during apoptosis ^28, 29^ but various forms of stress can also increase cytosolic levels of mtDNA ^30^. The architecture of mitochondrial cristae, the invaginations of the inner membrane into the matrix, appears to be a major regulator of mtDNA release ^31^. Because SMAC expression correlates with cytosolic mtDNA presence, SMAC may have a function in determining mitochondrial structure, which will affect spontaneous release of mtDNA. We found that SMAC-deficient MDA-MB-231 cells and MEFs had a different morphology of the mitochondrial network, with an increase in longer mitochondria and ellipticity index compared to control cells (Fig. 5A, B). MDA-MB-231 cells showed unaltered mitochondrial mass (Fig. S4A) but the mitochondrial membrane potential was reduced in SMAC-deficient cells (Fig. S4B). The oxygen consumption rate of SMAC-deficient MDA-MB-231 cells was unchanged (Fig. S4C).

**Figure 5:**
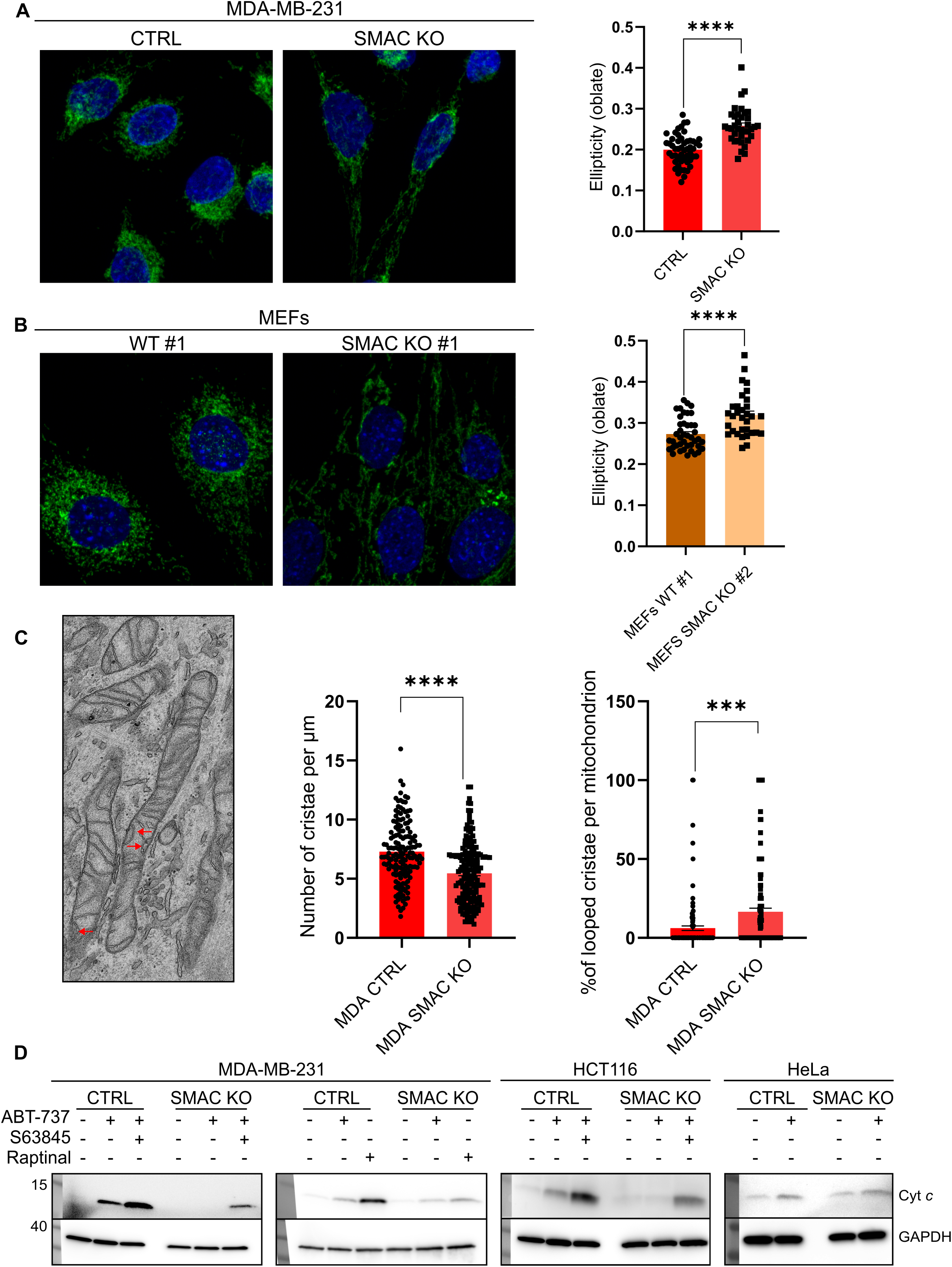
SMAC can regulate mitochondrial dynamics and integrity. **A**, **B**, morphology of the mitochondrial network in CTRL or SMAC-deficient MDA-MB-231 cells (A) and in wt or SMAC-deficient MEFs (B). Mitochondria were stained using anti-TOMM20 antibodies and observed by LSM880 Airyscan confocal microscopy. Each point in the graphs represents one cell from three independent experiments. Fusion was quantified using IMARIS software by segmenting mitochondria and measuring oblate ellipticity of the mitochondria. Means/SEM are shown. ****, p<0.0001 (unpaired t-test). **C**, morphology of the cristae in the mitochondria in CTRL or SMAC-deficient MDA-MB-231 cells observed using TEM. Red arrows show looped cristae. Cristae density and number of looped cristae were manually quantified. Means/SEM are shown. Each point indicates a mitochondrion from two independent experiments. ***, p<0.001. ****, p<0.0001 (unpaired t-test). **D**, cells from MDA-MB-231, HCT116 or HeLa cell lines were treated for 4 h with sub-lethal (2.5 µM ABT-737) or lethal (2.5 µM ABT-737 and 0.5 µM S63845) concentrations of BH3-mimetics. Raptinal was used at 1µM (4 h). Western blots are representative of three independent experiments.

When we visualized mitochondrial structure by electron microscopy, it became apparent that the cristae structure was different in SMAC-deficient cell. These cells had a significantly reduced cristae density (number of cristae along the mitochondria) and showed a pronounced increase in the number of ‘looped’ cristae in their mitochondria (Fig. 5C). The results suggest that SMAC deficiency leads to a structural defect in mitochondria, which is associated with a reduced spontaneous release of mtDNA. We tested whether SMAC deficiency also affected the apoptotic release of cytochrome *c*. Indeed, SMAC deficient MDA-MB-231 and HCT116 cells showed reduced release of cytochrome *c* upon permeabilization of the outer mitochondrial membrane, either by BH3-mimetics or by raptinal (Fig. 5D), which permeabilizes the outer mitochondrial membrane downstream of BAX/BAK ^32^. Curiously, no such effect was seen in HeLa cells (Fig. 5D).

### SMAC expression correlates with survival in cancer patients

Our results show regulation of sub-lethal caspase activation and of *in vitro* growth characteristics by SMAC in cancer cell lines. We have reported earlier that sub-lethal signals in the apoptosis pathway can drive metastasis in xenograft models and are associated with patient survival ^25^. Indeed, an association of enhanced SMAC expression and poor prognosis has been reported before in patients with metastatic melanoma ^33^. We analyzed the TCGA database for the association of SMAC expression and survival. As shown in Fig. 6, we found a significant association of high levels of SMAC mRNA expression and shorter patient survival in a number of cancer entities. The association was significant for lung adenocarcinoma and renal clear cell carcinoma across all stages, and mostly for the later tumor stages in the other tumor entities. Together with the functional data shown above, it seems most likely that the activation of sub-lethal signals downstream of mitochondria by SMAC translates into aggressive growth of cancer cells and thereby a poor prognosis of cancer patients.

**Figure 6:**
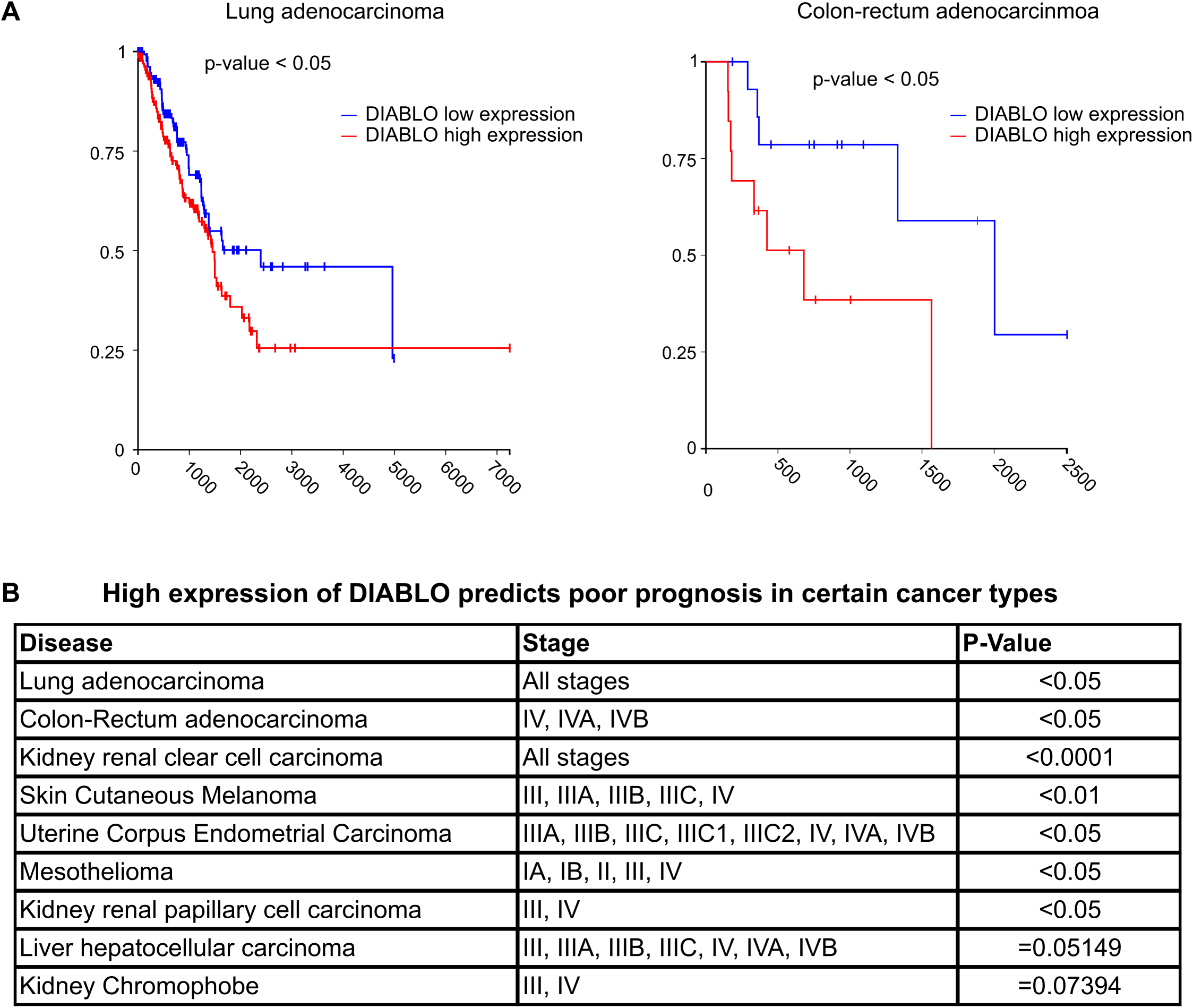
association of SMAC gene expression with prognosis in cancer patients. **A**, **B**, patient samples from TCGA were assigned according to the gene expression of DIABLO. **A**, Kaplan-Meier curves show the survival of patients in the upper and lower quartiles. **B**, various cancer entities from TCGA were stratified according to the DIABLO gene expression. High expression of DIABLO was associated with low survival probabilities in the indicated cancer types and stages (in each case, top quartile was compared to bottom quartile). Significance was estimated using log-rank test.

## Discussion

Our results indicate non-apoptotic functions of SMAC in both cytosol and mitochondria. In the cytosol, SMAC can regulate non-lethal caspase activity. SMAC also regulated mitochondrial morphology and integrity. The results extend the correlation of SMAC expression and tumor progression. One intriguing association is the level of SMAC with IFN signaling in cancer.

It was surprising to see the substantial difference regarding the requirement for SMAC between the tested cells. The observations are however in line with the results that have been reported for individual cell types. Notably, cells from DIABLO gene deficient mice showed no apoptosis defect; mostly analyzed were lymphocytes and thymocytes, which may be principally different, but embryonic stem cells and embryonic fibroblasts were also tested ^12^. On the other hand, human tumor cell lines near-completely depended on SMAC for caspase-activation following mitochondrial permeabilization ^13^. Because this could also be a mouse-human difference, we included a non-malignant human fibroblast line, which turned out also to have a reduced requirement for SMAC for caspase activation. We have no explanation of this differential response in molecular terms. The only obvious possibility would have been a difference in XIAP levels between the cells, but this was not found. It therefore appears on the basis of these data that SMAC dependency is found in transformed cells but not in non-malignant cells.

SMAC is one of several apoptosis signaling proteins whose expression levels are linked to cancer prognosis. It is clear that apoptosis is an anti-cancer mechanism: to become malignant, cells have to develop apoptosis resistance. The relationship is different however in established cancer. Here, high levels of pro-apoptotic proteins can be associated with poor prognosis – in addition to SMAC, this has been noted for BAX and BAK in melanoma – and high levels of BCL-2 are clearly associated with good prognosis in numerous cancer types ^34^. Because low activity in the apoptosis pathway will not only reduce apoptosis but also sub-lethal signals, these correlations may be the result of non-lethal activity in the apoptosis pathway. Sub-lethal signals can drive tumor growth ^35^, aggressive growth in cancer cell lines and metastasis in xenograft models ^25^. The results reported here show that SMAC can regulate the growth behavior of cancer cells. Part of this effect appears to be due to DNA damage and the DNA damage response, as a consequence of sub-lethal activity of the apoptosis pathway.

We have reported spontaneous activity of the mitochondrial apoptosis pathway in cancer cells and in mouse primary intestinal organoids before. The results shown here add the requirement for SMAC, presumably upon its release into the cytosol. The earlier results had suggested that mitochondrial ‘leakage’ occurs in unstimulated cells. Here we confirm this directly. The mechanism is not clear. Earlier studies of sub-lethal signaling have reported that individual fissioned-off mitochondria were particularly sensitive to BH3-mimetic but their permeabilization still required this stimulus ^22^. Both BAX and BAK appeared to be able to release SMAC and cytochrome *c* in healthy cells in culture. Double-deletion of BAX and BAK reduced but did not abolish the release. Loss of DRP1, on the other hand, appeared to block it completely; this may mean that indeed release occurs from mitochondria that have fissioned off from the network. Although the link between fission and apoptosis signaling is still incompletely understood, a role for DRP1 in mitochondrial permeabilization has been suggested long ago ^36^. DRP1 can directly interact with and activate BAX ^37^. What is still unclear is however how DRP1 could regulate mitochondrial permeabilization in the absence of BAX and BAK, as is suggested by the stronger phenotype of the DRP1-deficient cells (see above). Perhaps mitochondrial fission releases small amounts of intermembrane space content by simple leakage.

Our results suggest a role of SMAC in mitochondrial structure and integrity. SMAC-deficiency changed the morphology of mitochondria and reduced the release of cytochrome *c* upon MOMP. Most cytochrome *c* is not actually localized in the intermembrane space but within the cristae, which themselves are blocked off from the intermembrane space through cristae junctions ^38^. SMAC-dependent structural changes may further shift this balance and thereby reduce the cytochrome *c* available for release. Similarly, deletion of SMAC reduced the release of mtDNA into the cytosol. Because this release is also under the control of the cristae architecture ^31^, a regulatory effect of SMAC on cristae structure would have the same effect. Release of cytochrome *c* was also affected by the loss of SMAC. Because the greater part of cytochrome *c* (85 % in one estimate ^38^) is localized to the cristae, cristae structure may also affect the release of cytochrome *c*. The structural consequences of SMAC deficiency may therefore be the basis for changed mitochondrial integrity. The actual function of SMAC in mitochondria remains unresolved.

Cytosolic mtDNA is known to drive the IFN response through cGAS/STING stimulation ^39^. It seems a likely possibility that this SMAC-dependent regulation of the cytosolic levels of mtDNA is the basis for the correlation of IFN-induced genes and SMAC expression levels in cancer cells. cGAS stimulation in cancer cells affects the immune response to cancer through STING stimulation in infiltrating dendritic cells. In this way, SMAC is likely to be a co-determinant of the anti-cancer immune response, and this mechanism may be relevant to cancer progression and patient prognosis.

## Materials and Methods

### Cell lines and culture conditions

HeLa229 (HeLa) cells (ATCC Cat# CCL-2.1) and MDA-MB-231 cells (ATCC Cat# HTB-26) were cultured in RPMI 1640 medium (Thermo Fisher Scientific, Gibco) with 10% FCS (Sigma–Aldrich, #F7524). Mouse embryonic fibroblasts (MEFs) were generated from wild type and SMAC-deficient embryos and cultured in DMEM (Thermo Fisher Scientific, 41965062) with 10% FCS (Anprotec) and 50□µM 2-mercaptoethanol. WI-38 fibroblasts, a gift from Clemens Schmitt, Berlin, were cultured in DMEM supplemented with 10% FCS (Anprotec), 1% sodium pyruvate, 1% glutamax and 1% (v/v) penicillin/streptomycin (Gibco). The metastatic melanoma cell line 1205Lu (Meenhard Herlyn, Wistar Institute, Philadelphia) was cultured in TU2% melanoma medium containing 80% (v/v) MCDB153 (Sigma–Aldrich, #M7403), 20% (v/v) Leibovitz’s L-15, 2% (v/v) FCS (Thermo Fisher, Gibco), 5□µg/ml insulin (bovine, Sigma–Aldrich, #I4011), and 1.68□mM CaCl2. AGS cells (ATCC CRL-1739; obtained from ECACC, Sigma-Aldrich) were maintained in Ham’s F-12K medium (Gibco) with 10% FCS. HCT116 (from Christoph Borner), were cultured in DMEM with 10% FCS. Caco-2 cells (Tilman Brummer), were cultured in DMEM with 10% FCS. Jurkat cells (Georg Bornkamm, Munich) were cultured in DMEM with 10% FCS. All cells were cultured at 37□°C with 5% CO2. Gene-deficient cells were generated by CRISPR/Cas9 genome editing by transducing the cells with the lentiviral vector lentiCRISPR v2 (Addgene; Sanjana et al., 2014) and selection with puromycin (Invivogen). Guide RNAs used for CTRL CAD, SMAC, BAX, BAK and Caspase-9 as well as the BCL-X_L_-overexpressing cells have been described ^23, 24, 25, 40^. For generating Drp1-deficient cells, the gRNA AAATAGCTACGGTGAACCCG was used.

### Antibodies

The following primary antibodies were used: anti-γH2AX (#9718S, Cell Signaling), anti-GAPDH (MAB374, Millipore), anti-cleaved caspase-3 (#9661, Cell Signaling Technology), anti-caspase-3 (#9662, Cell Signaling Technology), anti-SMAC (#2954 and #15108, Cell Signaling Technology), anti-cytochrome c (#11940 and #12963, Cell Signaling Technology), anti-phospho-STING (#19781S, Cell Signaling Technology), anti-phospho-IRF3 (#ab76493, Abcam), anti-TFAM (#8076S, Cell Signaling Technology), anti-dsDNA (#ab27156, Abcam) and anti-TOM20 (#42406, Cell Signaling Technology). The following secondary antibodies were used: Alexa Fluor 488 donkey anti-rabbit (#711-545-152, Dianova), Alexa Fluor 488 donkey anti-mouse antibody (#115-545-062,Dianova), Alexa Fluor 647 donkey anti-rabbit (#711-605-152, Dianova), Cy5 donkey anti-mouse (#715-175-151, Dianova), goat IgG anti-mouse IgG (H+L)-HRP (#115-035-166, Dianova) and goat IgG anti-rabbit IgG (H+L)-HRP (#A6667, Sigma-Aldrich).

### Flow cytometry

Cells were trypsinized, combined with supernatants, centrifuged, and washed twice with PBS. They were re-suspended in PBS and fixed in 2% paraformaldehyde (PFA; Morphisto, catalog no. 11762.00500) for 15 minutes at room temperature. After washing with PBS containing 0.5% bovine serum albumin (BSA; PAN-Biotech, catalog no. P06-1391500), cells were further washed with DPBS containing 0.5% BSA and 0.5% saponin (Roth, catalog no. 4185.1), stained with an antibody against cleaved caspase-3 and analyzed using a FACSCalibur (Becton, Dickinson).

### Western Blot

Cells were lysed with RIPA buffer in the wells. Samples were mixed with Laemmli loading buffer and heated to 95□°C before loading to SDS PAGE. Nitrocellulose membranes were blocked with 5% milk. Proteins were detected with ECL substrate.

### Assay for anchorage-independent growth

Cells were plated in triplicate for each experiment. A 2% agarose solution was heated to 95□°C and combined with medium to prepare a 0.6% agarose basal layer. The cells were then mixed with medium and agarose to achieve a final concentration of 0.3% agarose in 6-well plates. The plates were kept at 37□°C, and 100□µl of cell culture medium was carefully added to each well twice a week. After an incubation period of 2–3 weeks, colonies were stained with a 0.005% crystal violet solution. Images of the plates were captured, and the stained colonies were quantified using ImageJ software.

### Migration Assay

For HeLa Cells, 5 × 10□cells were plated in a 6-well plate. Upon confluency, a 200 µl pipette tip was used to create a scratch in the monolayer. The wells were rinsed three times with PBS to remove any non-adherent cells. Medium containing 2% serum was then added, and images were captured immediately after scratching and twice daily in the same region of each well until the scratches had fully closed. The scratch areas were measured using ImageJ software and normalized to the scratch area of the control cell line at 0 hours. For MDA-MB-231 cells, the kit Radius™ Cell Migration Assay (#CBA-126, Cell BioLabs) was used according to manufacturer’s instructions. Briefly, 2.5 × 10□cells were plated in the provided 96-well plate. After attachment, the Radius™ Gel is removed and cells are allowed to migrate for 24 hours.

### Invasion Assay

The Falcon® Permeable Support for 24-well plates was used following the manufacturer’s instructions. Individual permeable supports (#353097, Corning) were placed into Falcon cell culture permeable support companion plates (#353504, Corning) and coated with 250 µg/ml Corning Matrigel basement membrane matrix (#354234, Corning) for 2 hours. Cells were seeded at a density of 5 × 10□ per condition and allowed to adhere overnight. After attachment, serum-free medium was added to the cells, while medium containing 10% FCS was added to the lower chamber of the well. The cells were incubated for 24 hours to allow invasion. Non-invasive cells were carefully removed using a wet cotton swab, and the membranes were fixed with absolute methanol before being stained with 1% crystal violet. Once dried, five representative images were captured for each permeable support at 10x magnification. The invaded cells were manually counted using ImageJ software.

### Caspase-3 activity assay

Cells from duplicates were pooled and lysed using buffer (#9803, Cell Signaling) supplemented with protease inhibitors (#04693132001, Roche). Ten microliters of the lysate were combined with reaction buffer (90 µl, MDB buffer) containing 11 µM Ac-DEVD-AMC (#I-1660.0005, Bachem), 100 µg/ml BSA, and 0.1% CHAPS, and the reaction was performed in triplicates. Analyses were conducted using a Spark 10M plate reader (Tecan). To precipitate active caspases, biotinylated VAD-fmk (#sc-311290A, Santa Cruz) was added 3 hours prior to cell harvesting. Cells were lysed with RIPA buffer, and aliquots of the lysate were heated to 95□°C in Laemmli buffer. Supernatants were incubated overnight with neutravidin beads (#29204, Thermo Fisher) at 4□°C. The beads were washed with RIPA buffer, re-suspended in Laemmli buffer, boiled, and analyzed via Western blotting to detect active caspase-3.

### RNA-sequencing

Library preparation for bulk-sequencing of poly(A)-RNA was done as described previously ^25^. Barcoded cDNA of each sample was generated with a Maxima RT polymerase (Thermo Fisher) using oligo-dT primer containing barcodes, unique molecular identifiers (UMIs) and an adaptor. 5’-Ends of the cDNAs were extended by a template switch oligo (TSO) and full-length cDNA was amplified with primers binding to the TSO-site and the adaptor. NEB UltraII FS kit was used to fragment cDNA. After end repair and A-tailing a TruSeq adapter was ligated and 3’-end-fragments were finally amplified using primers with Illumina P5 and P7 overhangs. The library was sequenced on a NextSeq 500 (Illumina) with 65 cycles for the cDNA in sread1 and 19 cycles for the barcodes and UMIs in read2. Data was processed using the published Drop-seq pipeline (v1.0) to generate sample- and gene-wise UMI tables ^41^. Reference genome (GRCh38) was used for alignment. Transcript and gene definitions were used according to GENCODE v38. Downstream analysis of feature counts was performed with R Studio. First, genes expressed below 10 read counts were excluded. Then, differentially expressed genes were determined using the R Bioconductor package DeSeq2 (version 1.30.1). Next, differentially expressed genes were filtered according to their adjusted p-value (<0.05) and absolute value of log2 fold change (>0.5). Heatmaps were generated using the R package pheatmap (version 1.0.12). Gene set enrichment analysis was performed using the R Bioconductor package fgsea (version 1.16.0) ^7^ together with the Hallmark pathways from the MSigDb. As suggested by fgsea, all differentially expressed genes were used for the analysis. Ggplot2 (version 3.3.5) package was used to create dot plots of pathway analysis.

### Immunofluorescence

Cells were incubated on IBIDI slides, fixed with 4% PFA for 10□min, followed by a 10□min permeabilization with PBS containing 0.2% Triton-X. Samples were blocked with PBS 5% BSA 0.2% Triton-X 0.1% Tween for 30 minutes. For mitochondrial DNA detection, samples were incubated with primary antibody anti-dsDNA 1:500 and anti-TOM20 1:400 in the blocking solution for an hour. Samples were washed then incubated with Alexa Flour 488 anti-mouse and Alexa Flour 647 anti-rabbit, both 1:500 in the blocking solution for an hour. For mitochondrial dynamics, samples were incubated with anti-TOM20 1:400 for an hour. Samples were washed and incubated with Alexa Flour 488 anti-rabbit for an hour. Samples were washed afterwards and stained with DAPI for 10 minutes. Samples were washed and covered with IBIDI mounting medium (#50001-4, IBIDI). Images were acquired using Zeiss LSM 880 with AiryScan. Analysis was done using ImageJ.

### Subcellular fractionation and mitochondrial DNA extraction

Cells were detached and re-suspended in DPBS. For detection of mitochondrial DNA in cytosol using qPCR, each sample was divided into two tubes. To extract the cytosol, cells were centrifuged and re-suspended in a solution of 70 mM Tris and 250 mM sucrose at pH 7. On ice, digitonin was added to 100 μg/ml for a minute followed by centrifugation at 920 g for 3 minutes. The supernatant was collected as cytosol and the pellet was solubilized in RIPA buffer for western blotting. To isolate mitochondrial DNA from the cytosolic fraction, QIAquick Nucleotide Removal Kit (#28306, QIAGEN) was used according the manufacturer’s protocol. Total DNA was isolated from whole cells with the DNeasy Blood & Tissue Kit (#69506, QIAGEN) according the manufacturer’s protocol. Equal volumes were used for the elution of DNA.

### qPCR

Equal volume of DNA extracted from cytosol or whole cell was mixed with SYBR Green master mix (#4472918, Applied Biosystems) and primers ^42^, followed by detection using QuantStudio 5 Real-Time PCR detection system (Applied Biosystems). Results were analyzed using QuantStudio Design and Analysis software v.1.5.2. The comparative CT method (ΔΔCT method) was used to quantify cytosolic mitochondrial D-loop region relative to *KCNJ* gene.

### Survival and correlation analysis

TCGA data were used for survival analysis assessing expression levels of the gene *DIABLO* and the pathological stage of tumors. Survival analysis of patients within highest and lowest quartiles of expression was done either with all tumor stages combined or with only late stages using the Xena platform ^43^. For correlation analysis, significantly downregulated interferon-regulated genes were identified using all available resources (in vitro/in vivo data of *Homo sapiens*) on INTERFEROME v2.0 database ^44^. The expression of significantly downregulated interferon-regulated genes was compared with expression levels of *DIABLO* in different tumors for significant correlation using Gene Expression Profiling Interactive Analysis (GEPIA) ^45^.

### Electron Microscopy

For transmission electron microscopy (TEM), samples were fixed using 2.5% glutaraldehyde (GA) in 0.1 M sodium cacodylate buffer (pH 7.4) at room temperature for one hour, followed by an overnight incubation at 4°C. After fixation, the samples were stored in 0.1 M sodium cacodylate buffer at 4°C until further processing. Contrasting was done with a single wash in 0.1 M sodium cacodylate buffer, followed by primary contrasting using a solution containing 1% osmium tetroxide (OsO_₄_) and 1.5% potassium ferrocyanide in the same buffer (1h on ice). The samples were washed twice with deionized water. A secondary contrasting step involved a 1% osmium tetroxide solution and was carried out on ice for one hour. Whole-mount staining was performed using a 1% uranyl acetate solution. For embedding, samples were dehydrated followed by a resin infiltration protocol using increasing concentrations of Durcupan in acetone (25%, 50%, 75%). Pure Durcupan infiltration followed (30 min at room temperature and an overnight incubation at 4°C). Final sample embedding was performed in Durcupan filled PCR tubes by polymerization for two days at 61°C in an oven. Ultrathin sections of 70 nm were cut and briefly stained with lead citrate for one minute. The imaging was done using a Talos L120C transmission electron microscope operated at 120 kV, with 10 cells imaged at a magnification of 13,500×.

## Acknowledgments

This work was supported by a grant from the Wilhelm Sander-Stiftung to G.H. (2019.019.1). We are grateful for support by the Core Facility for Electron Microscopy (EMcore) at the University Freiburg Medical Center—IMITATE is registered with the DFG (German Research Foundation) under the reference number RI_00555. We thank the staff of the Life Imaging Center (LIC) in the Hilde Mangold Haus (HMH), Albert-Ludwigs-University Freiburg for providing microscopy resources and support in image analysis. (LIC is funded by the Deutsche Forschungsgemeinschaft (DFG) – DFG 91b; Prof. Dr. G. Walz and Dr. R Nitschke DFG Project No. 317784314.

---

**Figure S1:**
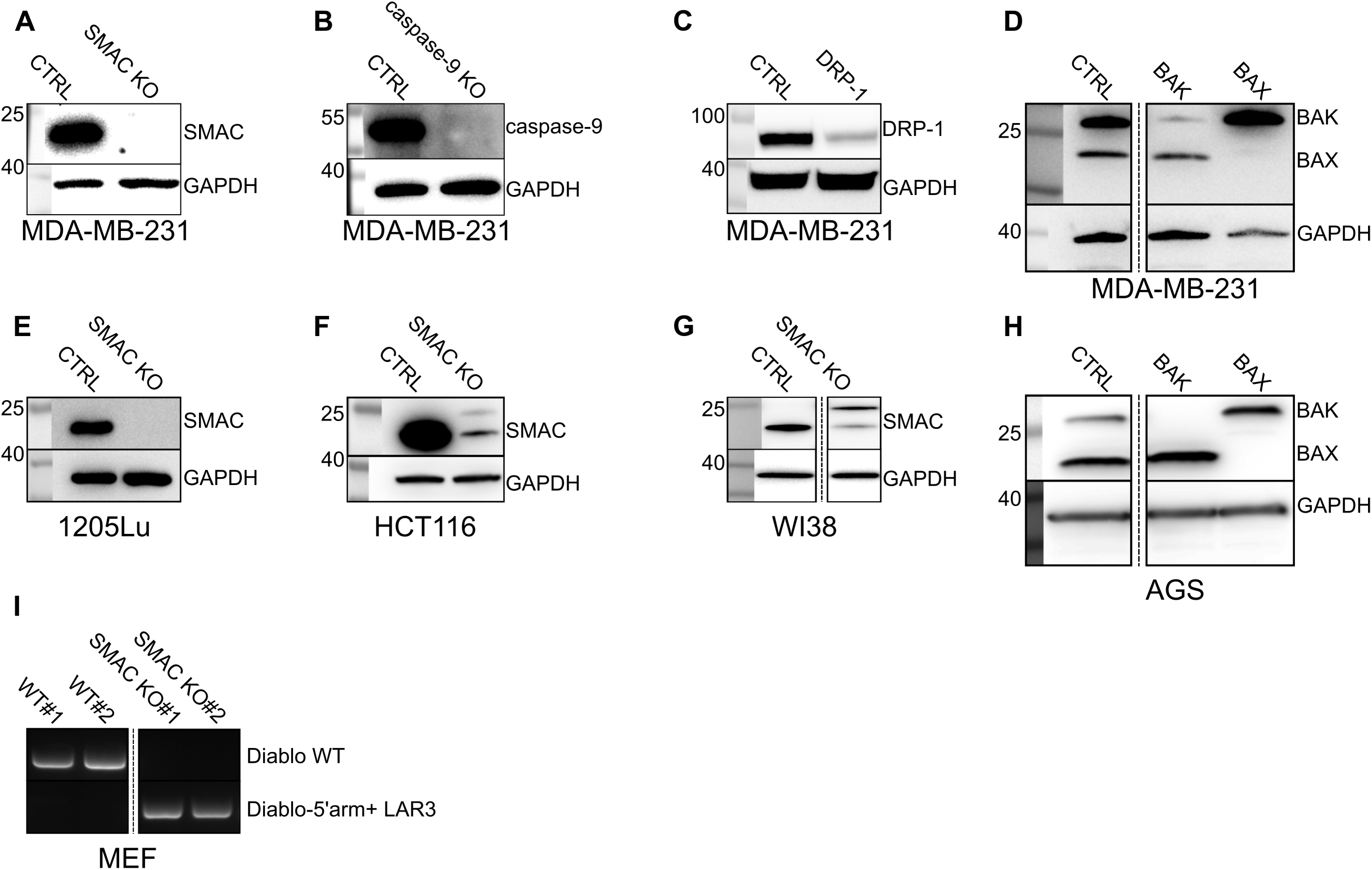
validation of gene deficiency in cell lines. **A-D**, deficiency of SMAC **(A)**, caspase-9 **(B)**, DRP-1 **(C)**, BAX or BAK **(D)** was validated in MDA-MB-231 by western blotting. **E-G**, western blot for 1205Lu **(E)**, HCT116 **(F)** and WI38 **(G)** cells validating SMAC deficiency. **H**, western blot for AGS cells validating BAX or BAK deficiency. **I**, Genotyping of the embryos from which MEF were isolated confirming successful gene editing.

**Figure S2:**
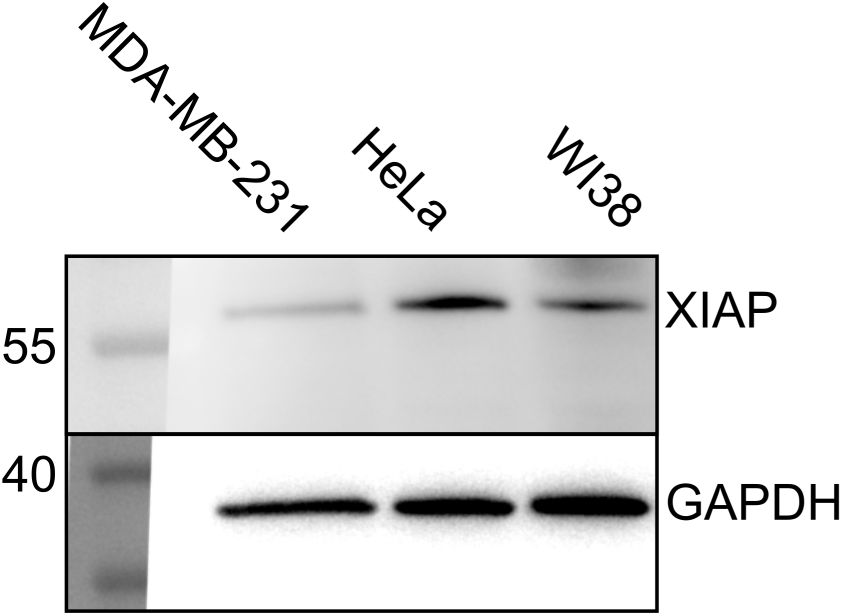
levels of XIAP in human cancer cells and primary cells. Levels of XIAP in MDA-MB-231, HeLa and WI38 cells were measured by western blotting. Shown is one of three independent experiments.

**Figure S3:**
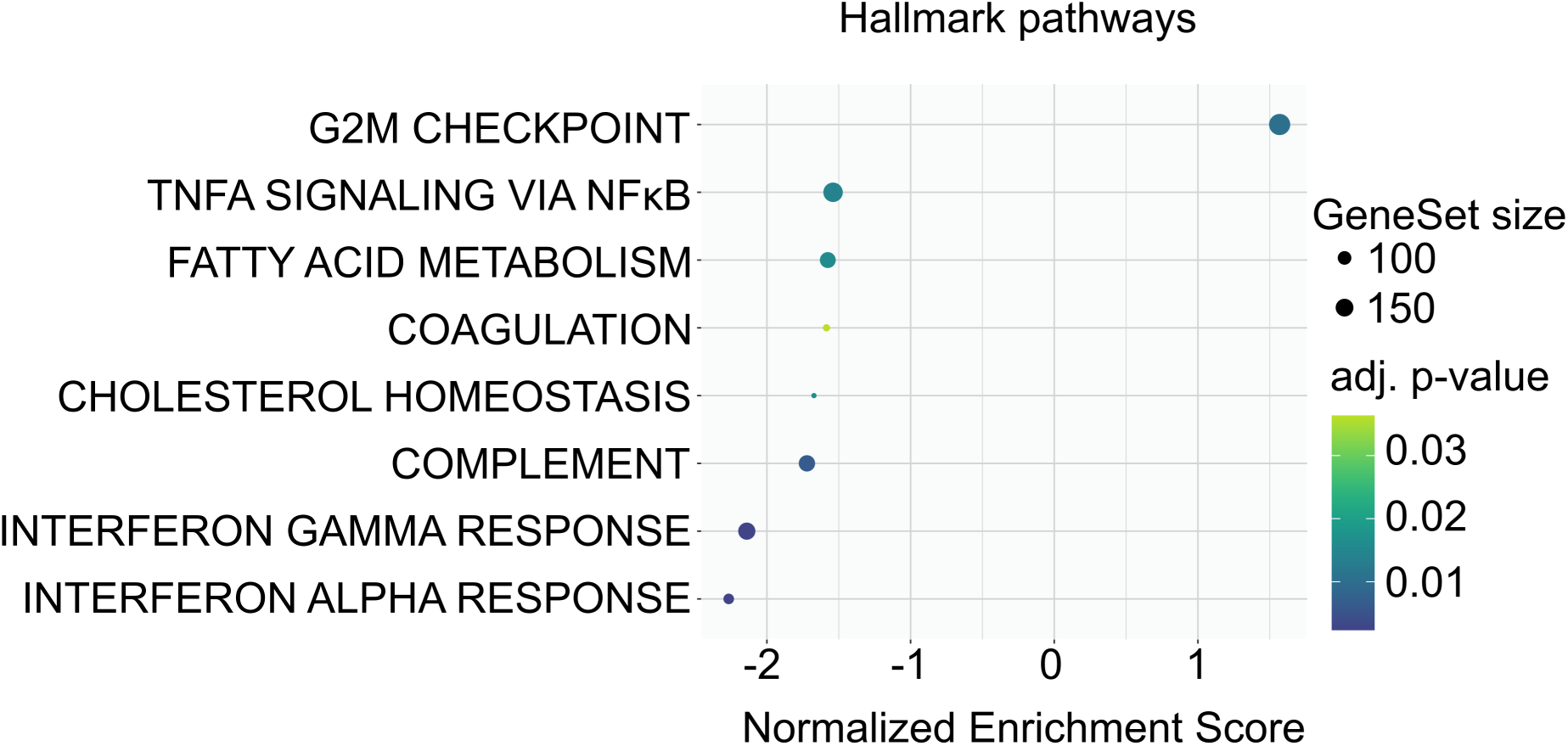
Pathway enrichment analysis in MDA-MB-231 SMAC KO. Enriched pathways in MDA-MB-231 were identified based on deferentially expressed genes in MDA-MB-231 SMAC KO cells based on hallmark pathways from the MSigDb.

**Figure S4:**
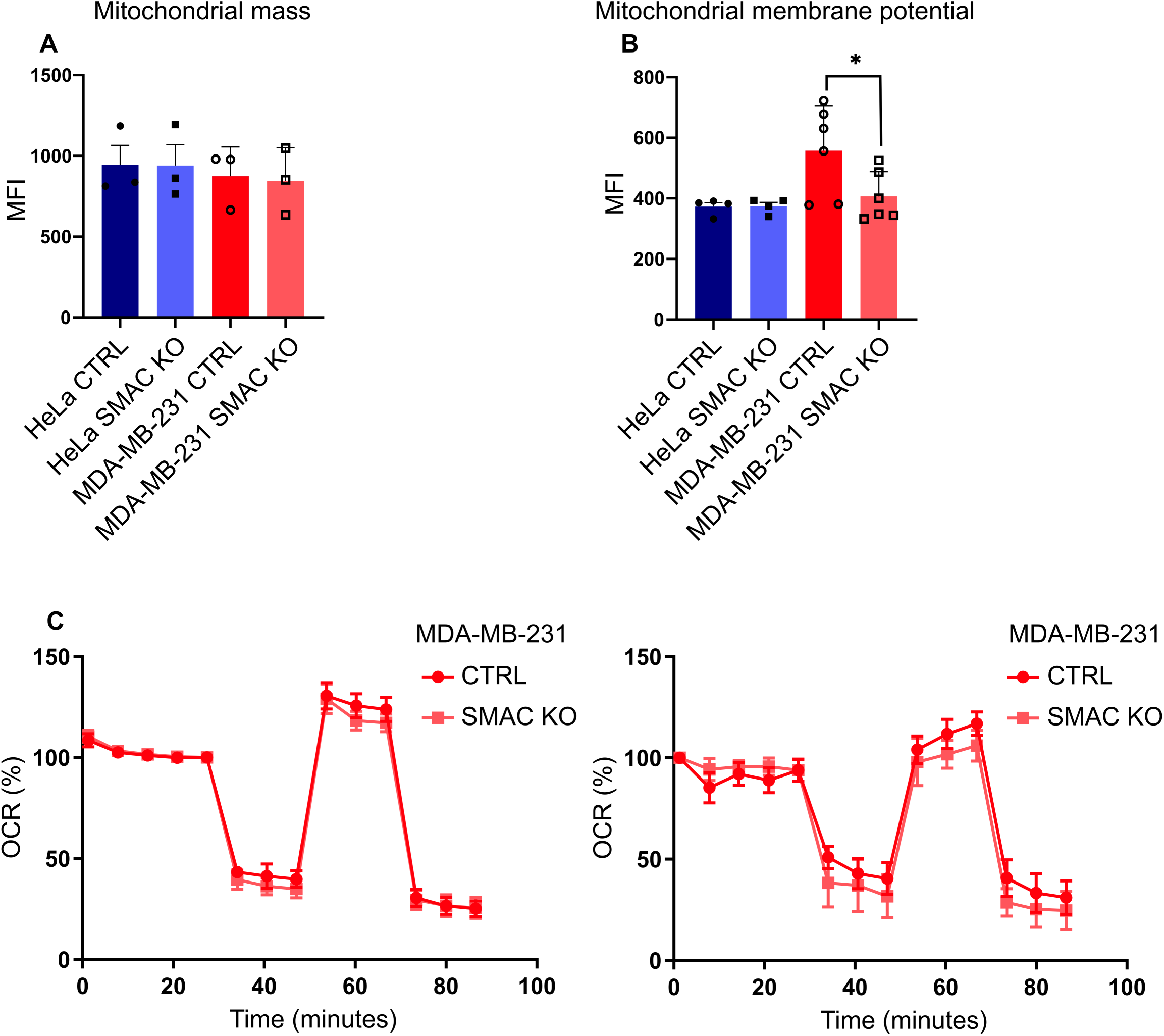
Mitochondrial changes in SMAC-deficient cells. **A**, mitochondrial mass was measured by flow cytometry using MitoTracker™ Green. Symbols represent independent experiments. **B**, membrane potential of HeLa and MDA-MB-231 cells was measured using MitoTracker™ Red CMXRos Dye by flow cytometry. Symbols represent independent experiments. **C**, oxygen consumption rate was measured in MDA-MB-231 cells using The Agilent Seahorse system. Two independent experiments are shown.

**Table S1:**
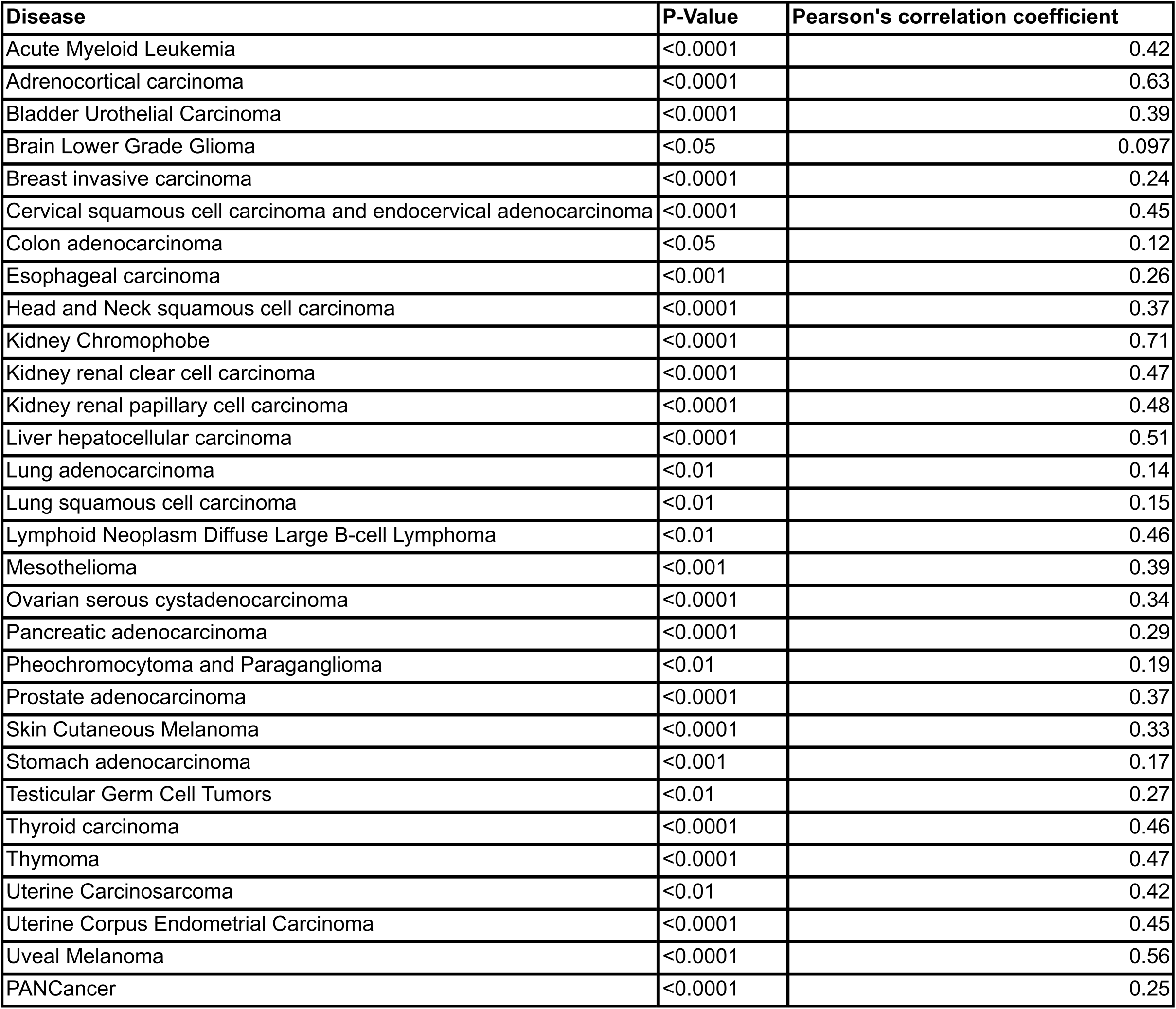
*Expression of DIABLO is positively correlated with the expression of interferon-regulated genes in multiple cancer* Pearson’s correlation coefficients and p-values are shown for the positive correlation between DIABLO and interferon-regulated genes extracted from RNA-sequencing data. Data are from The Cancer Genome Atlas.

| Disease | P-Value | Pearson's correlation coefficient |
| --- | --- | --- |
| Acute Myeloid Leukemia | <0.0001 | 0.42 |
| Adrenocortical carcinoma | <0.0001 | 0.63 |
| Bladder Urothelial Carcinoma | <0.0001 | 0.39 |
| Brain Lower Grade Glioma | <0.05 | 0.097 |
| Breast invasive carcinoma | <0.0001 | 0.24 |
| Cervical squamous cell carcinoma and endocervical adenocarcinoma | <0.0001 | 0.45 |
| Colon adenocarcinoma | <0.05 | 0.12 |
| Esophageal carcinoma | <0.001 | 0.26 |
| Head and Neck squamous cell carcinoma | <0.0001 | 0.37 |
| Kidney Chromophobe | <0.0001 | 0.71 |
| Kidney renal clear cell carcinoma | <0.0001 | 0.47 |
| Kidney renal papillary cell carcinoma | <0.0001 | 0.48 |
| Liver hepatocellular carcinoma | <0.0001 | 0.51 |
| Lung adenocarcinoma | <0.01 | 0.14 |
| Lung squamous cell carcinoma | <0.01 | 0.15 |
| Lymphoid Neoplasm Diffuse Large B-cell Lymphoma | <0.01 | 0.46 |
| Mesothelioma | <0.001 | 0.39 |
| Ovarian serous cystadenocarcinoma | <0.0001 | 0.34 |
| Pancreatic adenocarcinoma | <0.0001 | 0.29 |
| Pheochromocytoma and Paraganglioma | <0.01 | 0.19 |
| Prostate adenocarcinoma | <0.0001 | 0.37 |
| Skin Cutaneous Melanoma | <0.0001 | 0.33 |
| Stomach adenocarcinoma | <0.001 | 0.17 |
| Testicular Germ Cell Tumors | <0.01 | 0.27 |
| Thyroid carcinoma | <0.0001 | 0.46 |
| Thymoma | <0.0001 | 0.47 |
| Uterine Carcinosarcoma | <0.01 | 0.42 |
| Uterine Corpus Endometrial Carcinoma | <0.0001 | 0.45 |
| Uveal Melanoma | <0.0001 | 0.56 |
| PANCancer | <0.0001 | 0.25 |

## References

1. Glover HL, Schreiner A, Dewson G, Tait SWG. Mitochondria and cell death. Nature cell biology 2024.

2. Chipuk JE, Moldoveanu T, Llambi F, Parsons MJ, Green DR. The BCL-2 family reunion. Molecular cell 2010, 37(3): 299–310.

3. McArthur K, Kile BT. Apoptotic Caspases: Multiple or Mistaken Identities? Trends Cell Biol 2018.

4. Hao Z, Duncan GS, Chang CC, Elia A, Fang M, Wakeham A, et al. Specific ablation of the apoptotic functions of cytochrome C reveals a differential requirement for cytochrome C and Apaf-1 in apoptosis. Cell 2005, 121(4): 579–591.

5. Verhagen AM, Ekert PG, Pakusch M, Silke J, Connolly LM, Reid GE, et al. Identification of DIABLO, a mammalian protein that promotes apoptosis by binding to and antagonizing IAP proteins. Cell 2000, 102(1): 43–53.

6. Du C, Fang M, Li Y, Li L, Wang X. Smac, a mitochondrial protein that promotes cytochrome c-dependent caspase activation by eliminating IAP inhibition. Cell 2000, 102(1): 33–42.

7. Rehm M, Dussmann H, Prehn JH. Real-time single cell analysis of Smac/DIABLO release during apoptosis. The Journal of cell biology 2003, 162(6): 1031–1043.

8. Srinivasula SM, Hegde R, Saleh A, Datta P, Shiozaki E, Chai J, et al. A conserved XIAP-interaction motif in caspase-9 and Smac/DIABLO regulates caspase activity and apoptosis. Nature 2001, 410(6824): 112–116.

9. Zhang XD, Zhang XY, Gray CP, Nguyen T, Hersey P. Tumor necrosis factor-related apoptosis-inducing ligand-induced apoptosis of human melanoma is regulated by smac/DIABLO release from mitochondria. Cancer research 2001, 61(19): 7339–7348.

10. Rehm M, Huber HJ, Dussmann H, Prehn JH. Systems analysis of effector caspase activation and its control by X-linked inhibitor of apoptosis protein. The EMBO journal 2006, 25(18): 4338–4349.

11. McKenna S, Garcia-Gutierrez L, Matallanas D, Fey D. BAX and SMAC regulate bistable properties of the apoptotic caspase system. Scientific reports 2021, 11(1): 3272.

12. Okada H, Suh WK, Jin J, Woo M, Du C, Elia A, et al. Generation and characterization of Smac/DIABLO-deficient mice. Molecular and cellular biology 2002, 22(10): 3509–3517.

13. Saita S, Nolte H, Fiedler KU, Kashkar H, Venne AS, Zahedi RP, et al. PARL mediates Smac proteolytic maturation in mitochondria to promote apoptosis. Nature cell biology 2017, 19(4): 318–328.

14. Vince JE, Wong WW, Khan N, Feltham R, Chau D, Ahmed AU, et al. IAP antagonists target cIAP1 to induce TNFalpha-dependent apoptosis. Cell 2007, 131(4): 682–693.

15. Varfolomeev E, Blankenship JW, Wayson SM, Fedorova AV, Kayagaki N, Garg P, et al. IAP antagonists induce autoubiquitination of c-IAPs, NF-kappaB activation, and TNFalpha-dependent apoptosis. Cell 2007, 131(4): 669–681.

16. Silke J, Vucic D. IAP family of cell death and signaling regulators. Methods in enzymology 2014, 545: 35–65.

17. Paul A, Krelin Y, Arif T, Jeger R, Shoshan-Barmatz V. A New Role for the Mitochondrial Pro-apoptotic Protein SMAC/Diablo in Phospholipid Synthesis Associated with Tumorigenesis. Mol Ther 2018, 26(3): 680–694.

18. Pandey SK, Shteinfer-Kuzmine A, Chalifa-Caspi V, Shoshan-Barmatz V. Non-apoptotic activity of the mitochondrial protein SMAC/Diablo in lung cancer: Novel target to disrupt survival, inflammation, and immunosuppression. Front Oncol 2022, 12: 992260.

19. Shoshan-Barmatz V, Arif T, Shteinfer-Kuzmine A. Apoptotic proteins with non-apoptotic activity: expression and function in cancer. Apoptosis 2023, 28(5-6): 730–753.

20. Goldstein JC, Waterhouse NJ, Juin P, Evan GI, Green DR. The coordinate release of cytochrome c during apoptosis is rapid, complete and kinetically invariant. Nature cell biology 2000, 2(3): 156–162.

21. Ichim G, Lopez J, Ahmed SU, Muthalagu N, Giampazolias E, Delgado ME, et al. Limited mitochondrial permeabilization causes DNA damage and genomic instability in the absence of cell death. Molecular cell 2015, 57(5): 860–872.

22. Cao K, Riley JS, Heilig R, Montes-Gomez AE, Vringer E, Berthenet K, et al. Mitochondrial dynamics regulate genome stability via control of caspase-dependent DNA damage. Developmental cell 2022.

23. Brokatzky D, Dorflinger B, Haimovici A, Weber A, Kirschnek S, Vier J, et al. A non-death function of the mitochondrial apoptosis apparatus in immunity. The EMBO journal 2019, 38(11): e102325.

24. Dorflinger B, Badr MT, Haimovici A, Fischer L, Vier J, Metz A, et al. Mitochondria supply sub-lethal signals for cytokine secretion and DNA-damage in H. pylori infection. Cell death and differentiation 2022.

25. Haimovici A, Hofer C, Badr MT, Bavafaye Haghighi E, Amer T, Boerries M, et al. Spontaneous activity of the mitochondrial apoptosis pathway drives chromosomal defects, the appearance of micronuclei and cancer metastasis through the Caspase-Activated DNAse. Cell death & disease 2022, 13(4): 315.

26. West AP, Khoury-Hanold W, Staron M, Tal MC, Pineda CM, Lang SM, et al. Mitochondrial DNA stress primes the antiviral innate immune response. Nature 2015, 520(7548): 553–557.

27. Hong C, Schubert M, Tijhuis AE, Requesens M, Roorda M, van den Brink A, et al. cGAS-STING drives the IL-6-dependent survival of chromosomally instable cancers. Nature 2022, 607(7918): 366–373.

28. McArthur K, Whitehead LW, Heddleston JM, Li L, Padman BS, Oorschot V, et al. BAK/BAX macropores facilitate mitochondrial herniation and mtDNA efflux during apoptosis. Science 2018, 359(6378).

29. Riley JS, Quarato G, Cloix C, Lopez J, O’Prey J, Pearson M, et al. Mitochondrial inner membrane permeabilisation enables mtDNA release during apoptosis. The EMBO journal 2018.

30. Hu MM, Shu HB. Mitochondrial DNA-triggered innate immune response: mechanisms and diseases. Cell Mol Immunol 2023, 20(12): 1403–1412.

31. He B, Yu H, Liu S, Wan H, Fu S, Liu S, et al. Mitochondrial cristae architecture protects against mtDNA release and inflammation. Cell reports 2022, 41(10): 111774.

32. Palchaudhuri R, Lambrecht MJ, Botham RC, Partlow KC, van Ham TJ, Putt KS, et al. A Small Molecule that Induces Intrinsic Pathway Apoptosis with Unparalleled Speed. Cell reports 2015, 13(9): 2027–2036.

33. Gutta C, Rahman A, Aura C, Dynoodt P, Charles EM, Hirschenhahn E, et al. Low expression of pro-apoptotic proteins Bax, Bak and Smac indicates prolonged progression-free survival in chemotherapy-treated metastatic melanoma. Cell death & disease 2020, 11(2): 124.

34. Labi V, Erlacher M. How cell death shapes cancer. Cell death & disease 2015, 6: e1675.

35. Liu X, Li F, Huang Q, Zhang Z, Zhou L, Deng Y, et al. Self-inflicted DNA double-strand breaks sustain tumorigenicity and stemness of cancer cells. Cell research 2017, 27(6): 764–783.

36. Karbowski M, Lee YJ, Gaume B, Jeong SY, Frank S, Nechushtan A, et al. Spatial and temporal association of Bax with mitochondrial fission sites, Drp1, and Mfn2 during apoptosis. The Journal of cell biology 2002, 159(6): 931–938.

37. Jenner A, Pena-Blanco A, Salvador-Gallego R, Ugarte-Uribe B, Zollo C, Ganief T, et al. DRP1 interacts directly with BAX to induce its activation and apoptosis. The EMBO journal 2022, 41(8): e108587.

38. Scorrano L, Ashiya M, Buttle K, Weiler S, Oakes SA, Mannella CA, et al. A distinct pathway remodels mitochondrial cristae and mobilizes cytochrome c during apoptosis. Developmental cell 2002, 2(1): 55–67.

39. West AP, Shadel GS. Mitochondrial DNA in innate immune responses and inflammatory pathology. Nature reviews Immunology 2017, 17(6): 363–375.

40. Gradzka-Boberda S, Gentle IE, Hacker G. Pattern Recognition Receptors of Nucleic Acids Can Cause Sublethal Activation of the Mitochondrial Apoptosis Pathway during Viral Infection. Journal of virology 2022: e0121222.

41. Macosko EZ, Basu A, Satija R, Nemesh J, Shekhar K, Goldman M, et al. Highly Parallel Genome-wide Expression Profiling of Individual Cells Using Nanoliter Droplets. Cell 2015, 161(5): 1202–1214.

42. Bryant JD, Lei Y, VanPortfliet JJ, Winters AD, West AP. Assessing Mitochondrial DNA Release into the Cytosol and Subsequent Activation of Innate Immune-related Pathways in Mammalian Cells. Curr Protoc 2022, 2(2): e372.

43. Goldman MJ, Craft B, Hastie M, Repecka K, McDade F, Kamath A, et al. Visualizing and interpreting cancer genomics data via the Xena platform. Nature biotechnology 2020, 38(6): 675–678.

44. Rusinova I, Forster S, Yu S, Kannan A, Masse M, Cumming H, et al. Interferome v2.0: an updated database of annotated interferon-regulated genes. Nucleic acids research 2013, 41(Database issue): D1040–1046.

45. Tang Z, Kang B, Li C, Chen T, Zhang Z. GEPIA2: an enhanced web server for large-scale expression profiling and interactive analysis. Nucleic acids research 2019, 47(W1): W556–W560.

